# Bioinformatics Analysis Suggests That SE_1780 protein from Staphylococcus epidermidis may be a member of the Fph Family of Lipases

**DOI:** 10.1101/2024.10.06.616924

**Authors:** Maya Qaddourah, Sajith Jayasinghe

**Affiliations:** Department of Chemistry and Biochemistry, California State University, San Marcos, CA 92096

## Abstract

The Protein Databank entry for protein SE_1780 from Staphylococcus epidermidis lists the function as unknown. We leveraged the framework outlined in the Biochemistry Authentic Scientific Inquiry Laboratory and used bioinformatics tools to ascertain the function of the protein. Based on our analysis we posit that SE_1780 is a lipase of the **α**/**β** hydrolase family with a proposed active site catalytic triad composed of Ser144, Asp235, and His269. Further we identified the lipase FphD as having significant sequence identity to protein SE 1780, and suggest that the protein is a member of the Fph family of lipases from S. epidermidis.

## Introduction

Protein SE_1780 from Staphylococcus epidermidis (the POI) contains 289 residues and is composed of a central **β**-sheet surrounded by a collection of alpha helices (Figure 1) ^1^. The beta sheet is comprised of a total of eight **β**-strands, with seven strands arranged parallel to each other, while the remaining strand (**β**3), located at the leading edge of the sheet, runs anti-parallel to the rest. Of the seven **α**-helices five are located on one face of the sheet and two on the other. This core structure is similar to the architecture of enzymes containing the **α**/**β**-hydrolase fold ^2^. The PDB entry for the protein, 3FLE, lists the function as unknown. We leveraged the framework described in the Biochemistry Authentic Scientific Inquiry Laboratory (BASIL)^3,4^ Course Based Undergraduate Research Experience (CURE) and carried out a bioinformatics analysis to hypothesize a possible function for the POI.

**Figure 1.**
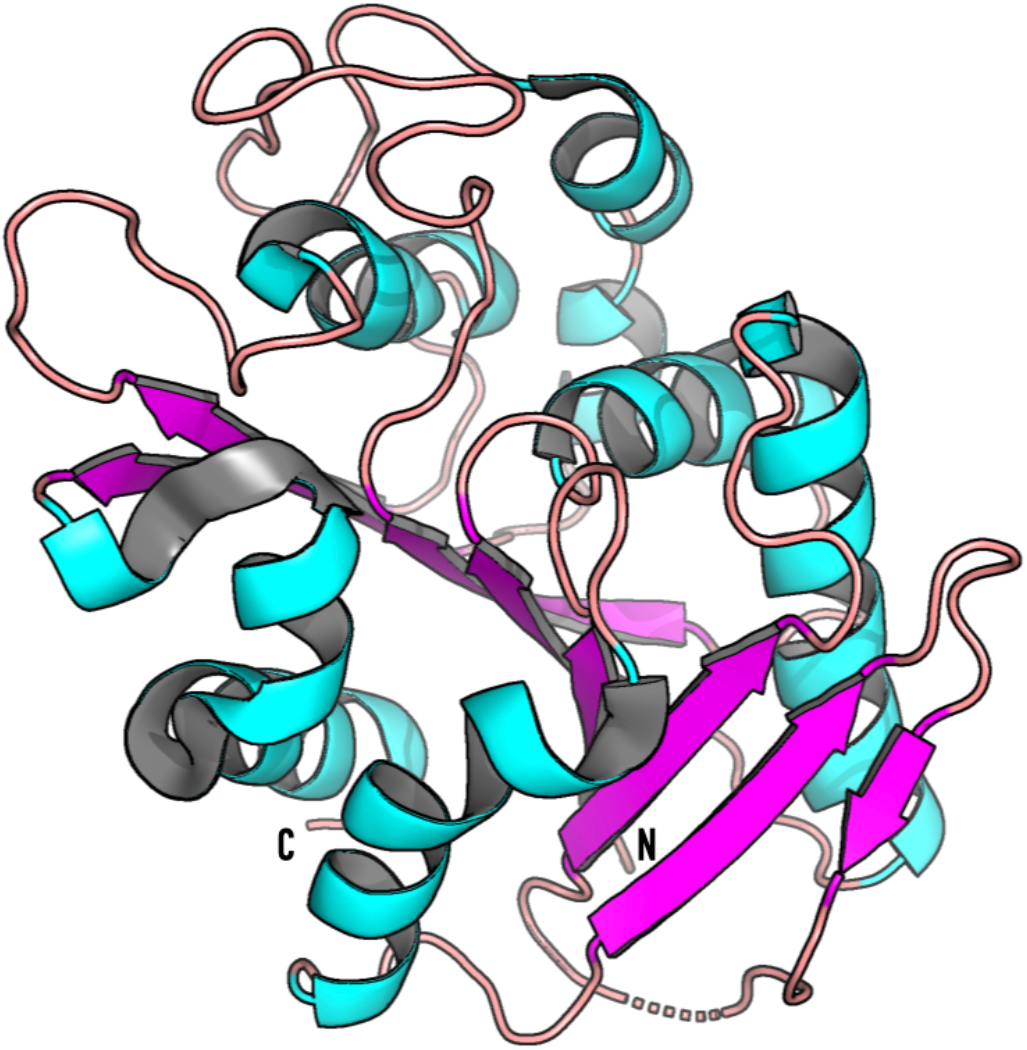
Cartoon representation of the structure of S. epidermidis (PDB 3FLE). N, and C indicate the N- and C-termini respectively. The protein contains a central beta-sheet composed of eight beta-stands with the leading-edge **β**-strand running antiparallel to the rest. A total of seven alpha helices are distributed on either side of the sheet.

## Materials and Methods

*Structural comparison of the POI with proteins in the Protein Data Bank (PDB) were conducted using the DALI server (http://ekhidna2.biocenter.helsinki.fi/dali/). DALI results with the highest Z-scores were used as structural homologs. BLAST search was carried out using the NCBI BLAST server (https://blast.ncbi.nlm.nih.gov/Blast.cgi) and pairwise sequence alignments were carried out using EMBOSS Needle (https://www.ebi.ac.uk/jdispatcher/psa/emboss_needle). Molecular visualization and graphics were created using PyMol (The PyMOL Molecular Graphics System, Version 3.0 Schrödinger, LLC)*.

## Results and Discussion

### Structural Similarity Suggests that Protein SE_1780 from Staphylococcus epidermidis is a Lipase

To gain insights as to the function of the SE_1780 protein we compared the POI to proteins with known structure using the protein structure comparison server DALI ^5^. The server identified several PDB entries with z-scores greater than 4 with sequence identities ranging from 7-30% (Table S1). The result with the highest Z-score (31.5), highest sequence identity (30%), and lowest RMSD (1.7) to the POI was the protein product from gene lin2722 from Listeria innocua (PDB ID: 3DS8). Unfortunately, the PDB lists the function of this protein as unknown and was thus unable to shed light on the function of our POI. However, a literature search found a report of a computational and in-vitro analysis suggesting that the protein from gene lin2722 is an **α**/**β**-hydrolase ^6^. The structure of the POI is also similar to the structures of lipase Lip_vut1 from the goat rumen metagenome (PDB ID 6NKC), Bacillus subtilis lipase A (PDB ID: 5CT6), and Bacillus pumilus Lipase A (PDB ID: 7R1K) with sequence identities of 24%, 20%, and 20% respectively (Table S1). Many of the other results of the DALI search were PDB entries of Bacillus subtilis lipase A crystalized under various conditions (for example PDB IDs 1R50 and 1I6W) but the results also contained entries described as carboxyesterases, arylesterases, and phospholipases. Thus, the DALI search indicates the possibility that our POI is a lipase in the **α**/**β** hydrolase family.

### The POI may be a homolog of S. aureus FphD

A BLAST search against the proteins in the Esterase and **α**/**β**-hydrolase enzymes and relatives (ESTHER) database ^7^ indicated that the POI has a 64% sequence identity to the Staphylococcus aureus cell surface protein FphD. FphD is yet to be functionally and structurally characterized but is described as belonging to a group of ten **α**/**β**-hydrolase enzymes known as fluorophosphonate-binding hydrolases (Fphs) which are produced during biofilm formation^8–10^. Three other members of this family of proteins, FphB FphF , FphH have been functionally characterized as lipases that cleave lipid ester substrates, and FphF and FphH have been structurally characterized^9,11^.

### The Catalytic Triad of Protein Se_1780 is comprised of Ser144, Asp 235, and His 269

To ascertain the nature of the active site we compared the structure of our POI with that of Bacillus subtilis lipase A (Figure 2) ^12^. The POI exhibits significant structural similarity with B. subtilis Lipase A (rmsd 2.39). We selected the subtilis lipase A instead of any of the Fph proteins based on the lower rmsd observed between subtilis Lipase A and the POI. The main structural differences between B. subtilis lipase A and the POI are in the beta sheet and the loop between **β**-stands 6 and 7. Bacillus subtilis lipase A is 77 residues shorter than the POI and its core beta-sheet contains only six strands, as opposed to the 8 in the POI. B. subtilis lipase A also has a much shorter loop between **β**-strands 6 and 7 compared to the POI (Figure 2).

**Figure 2.**
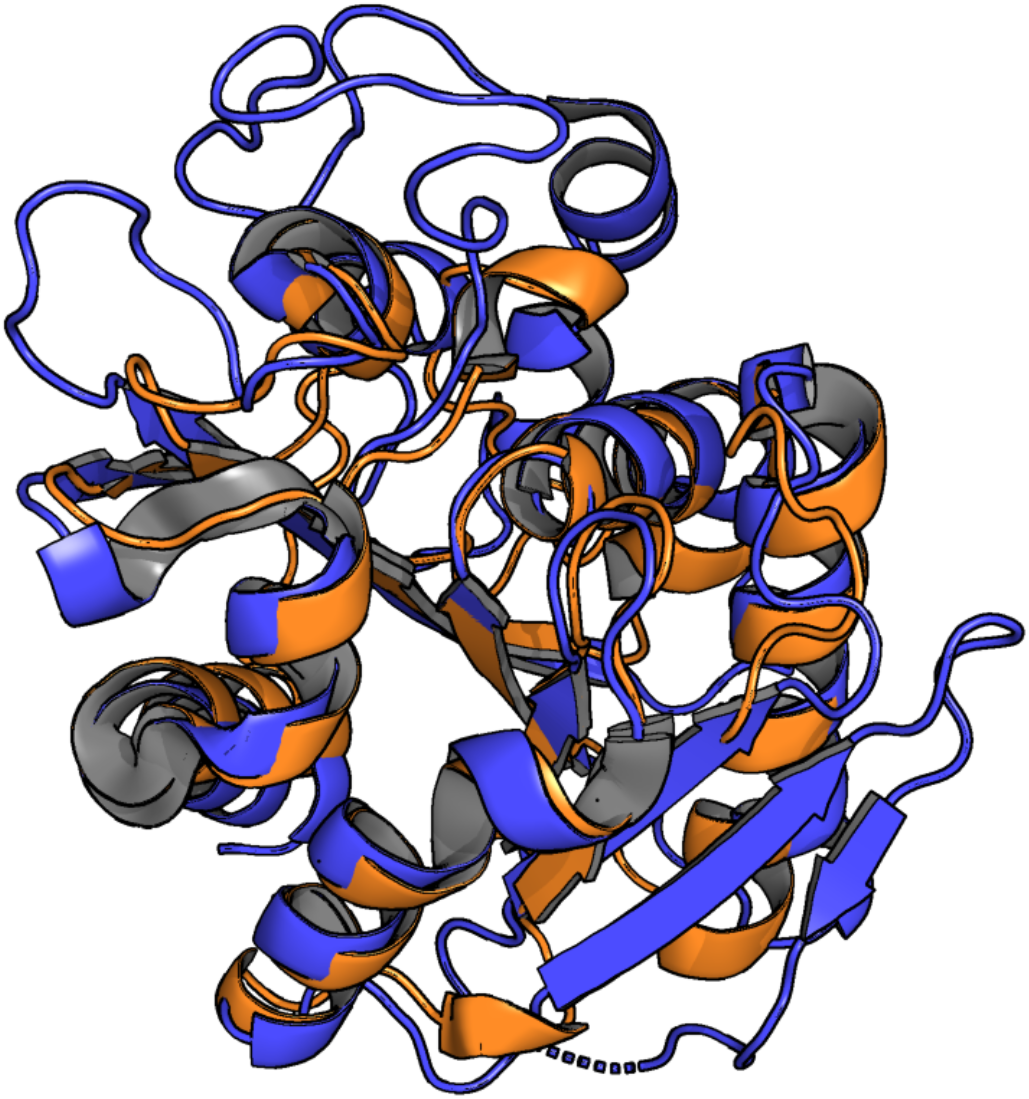
Comparison of crystal structure of the POI (blue) with that of B. Subtilis Lipase A (PDBID:1I6W, orange). The POI has significant structural similarity to subtilis Lipase A but differs in two main aspects. The POI is 77 residues longer than subtilis Lipase A and contains eight beta strands in its core beta sheet compared to six in the Lipase A. The POI has a much longer loop between beta-stands 6 and 7 compared to Lipase A.

The region of the POI that correspond to the location of the catalytic triad residues (Ser77, Asp133, and His 156) of B. subtilis show significant structural similarity, and therefore we identified the putative catalytic triad of the POI as Ser144, Asp235, and His269 (Figure 3). The proposed nucleophilic Ser144 is located on the loop between strand **β**5 and **β**6, Asp 235 is located at the end of strand **β**7, and the His269 is situated in the loop after strand **β**8. The catalytic serine is found within the sequence GHSMG which conforms to the conserved GXSXG peptapeptide sequence motif found in lipolytic enzymes ^13^. Based on the analysis of diverse lipases it has been suggested that variations in the conserved pentapeptide sequence may be used to categorize lipases into 19 families. The specific pentapeptide sequence, GHSMG, present in the POI has not been previously identified ^13^. The same pentapeptide sequence is present in FphD but not in the other Fph proteins.

**Figure 3.**
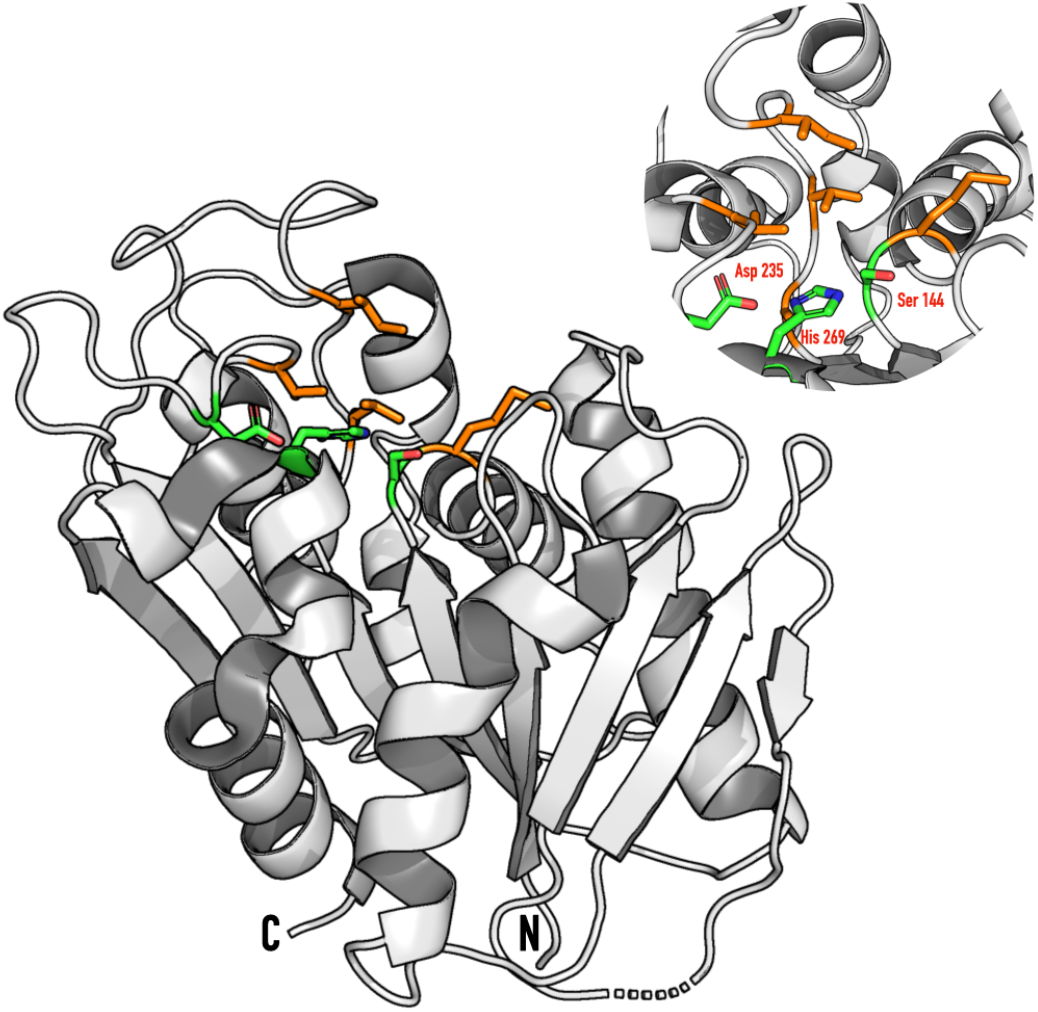
The proposed catalytic triad of the POI. Based on the location of the catalytic triad of subtilis Lipase A we propose that the catalytic triad of the POI is comprised of residues Ser144, Asp 235, and His269 (inset). The proposed nucleophilic Ser 144 is located within the sequence GHSMG which conforms to the GXSXG peptapeptide sequence (where X denotes any amino acids) motif observed in lipolytic enzymes. The active site is surface exposed and lined with hydrophobic residues (orange, Met 145, Ala 173, Val 175, Ile 179, and Val 238).

Give the hydrophobic nature of their substrates, lipases have active site cavities lined with hydrophobic side chains. We find that the catalytic triad of the POI is surrounded by several hydrophobic residues: Met 145, Ala 173, Val 175, Ile 179, and Val 238. In many lipases the hydrophobic active site is covered by a ‘lid’, composed of an amphipathic helix, that protects the active site ^14^. The active site of the POI is surface exposed and is not covered by a lid region, and in this regard is similar to lipases, such as B. subtilis lipase A, that also does not have a lid.

#### Conclusion

Using the BASIL CURE framework we postulate that protein SE_1780 from Staphylococcus epidermidis identified in PDB entry 3FLE is a lipase and that the protein may be member of the recently identified fluorophosphonate-binding hydrolases. We are in the process of expressing and purifying the protein to validate our hypothesis.

---

**Supplementary Table 1.**
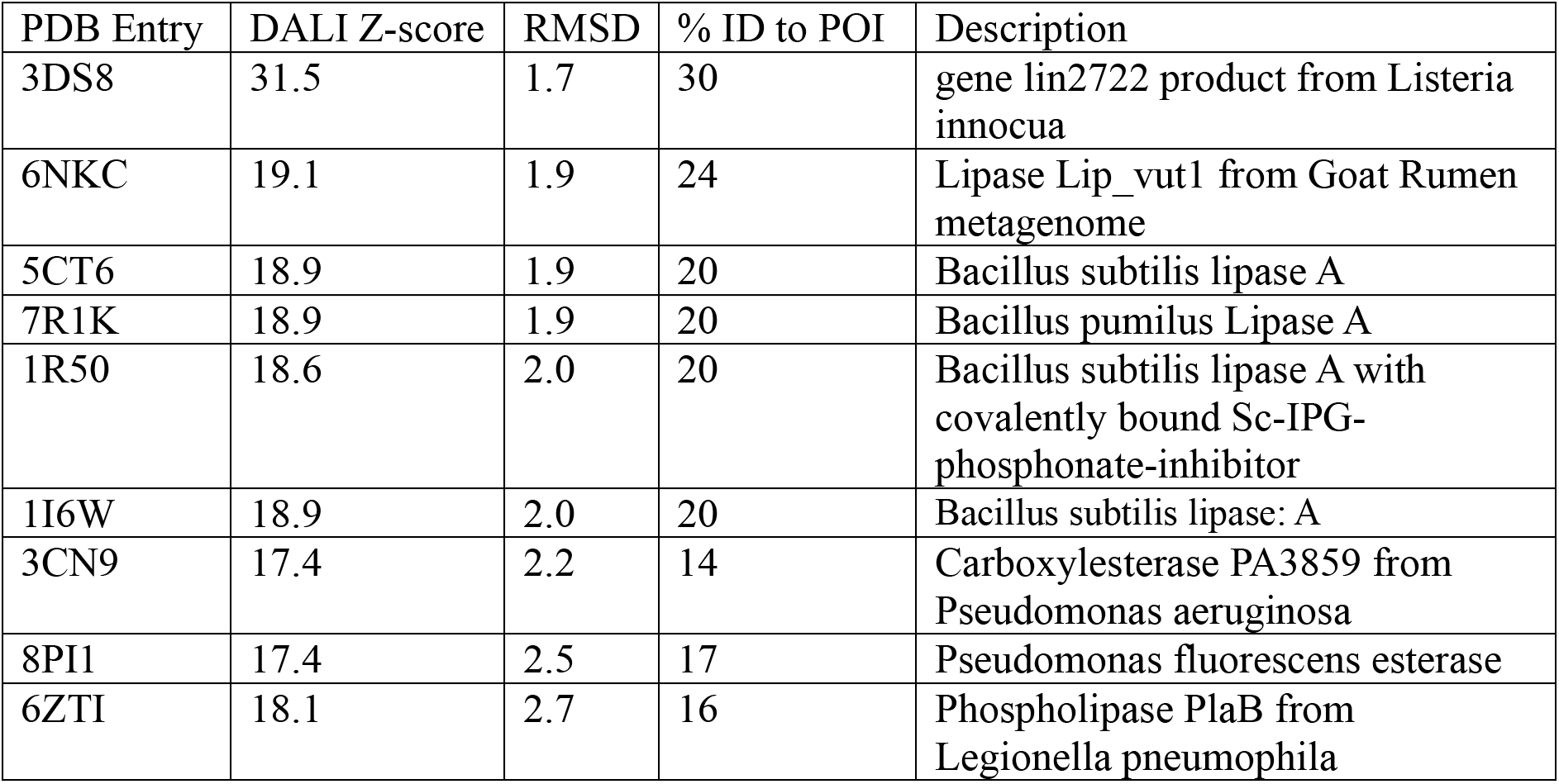

| PDB Entry | DALI Z-score | RMSD | % ID to POI | Description |
| --- | --- | --- | --- | --- |
| 3DS8 | 31.5 | 1.7 | 30 | gene lin2722 product from *Listeria innocua* |
| 6NKC | 19.1 | 1.9 | 24 | Lipase Lip_vut1 from Goat Rumen metagenome |
| 5CT6 | 18.9 | 1.9 | 20 | *Bacillus subtilis* lipase A |
| 7R1K | 18.9 | 1.9 | 20 | *Bacillus pumilus* Lipase A |
| 1R50 | 18.6 | 2.0 | 20 | *Bacillus subtilis* lipase A with covalently bound Sc-IPG-phosphonate-inhibitor |
| 1I6W | 18.9 | 2.0 | 20 | *Bacillus subtilis* lipase: A |
| 3CN9 | 17.4 | 2.2 | 14 | Carboxylesterase PA3859 from *Pseudomonas aeruginosa* |
| 8PI1 | 17.4 | 2.5 | 17 | *Pseudomonas fluorescens* esterase |
| 6ZTI | 18.1 | 2.7 | 16 | Phospholipase PlaB from *Legionella pneumophila* |

## References

(1) 3FLE RCSB Entry. 10.2210/pdb3FLE/pdb.

(2) Nardini, M.; Dijkstra, B. W. α/β Hydrolase Fold Enzymes: The Family Keeps Growing. Curr. Opin. Struct. Biol. 1999, 9 (6), 732–737. 10.1016/s0959-440x(99)00037-8.

(3) Roberts, R.; Hall, B.; Daubner, C.; Goodman, A.; Pikaart, M.; Sikora, A.; Craig, P. Flexible Implementation of the BASIL CURE. Biochem. Mol. Biol. Educ. 2019, 47 (5), 498–505. 10.1002/bmb.21287.

(4) Koeppe, J. R.; Roberts, R.; Hall, B. L.; Craig, P. A. The BASIL Cure: Using Structure to Predict Function in Protein Biochemistry. Biophys. J. 2023, 122 (3), 297a–298a. 10.1016/j.bpj.2022.11.1681.

(5) Holm, L. Using Dali for Protein Structure Comparison. Methods Mol. Biol. (Clifton, NJ) 2020, 2112, 29–42. 10.1007/978-1-0716-0270-6_3.

(6) Sharkawy, M.; Carter, A. A.; Craig, P. Function Identification of the Protein Product of Gene Lin2722 from Listeria Innocua Using Computational and In-Vitro Techniques. Biophys. J. 2019, 116 (3), 67a. 10.1016/j.bpj.2018.11.406.

(7) Lenfant, N.; Hotelier, T.; Velluet, E.; Bourne, Y.; Marchot, P.; Chatonnet, A. ESTHER, the Database of the α/β-Hydrolase Fold Superfamily of Proteins: Tools to Explore Diversity of Functions. Nucleic Acids Res. 2012, 41 (D1), D423–D429. 10.1093/nar/gks1154.

(8) Lentz, C. S.; Sheldon, J. R.; Crawford, L. A.; Cooper, R.; Garland, M.; Amieva, M. R.; Weerapana, E.; Skaar, E. P.; Bogyo, M. Identification of a S. Aureus Virulence Factor by Activity-Based Protein Profiling (ABPP). Nat. Chem. Biol. 2018, 14 (6), 609–617. 10.1038/s41589-018-0060-1.

(9) Fellner, M. Newly Discovered Staphylococcus Aureus Serine Hydrolase Probe and Drug Targets. ADMET DMPK 2021, 10 (2), 107–114. 10.5599/admet.1137.

(10) Fellner, M.; Lentz, C. S.; Jamieson, S. A.; Brewster, J. L.; Chen, L.; Bogyo, M.; Mace, P. D. Structural Basis for the Inhibitor and Substrate Specificity of the Unique Fph Serine Hydrolases of Staphylococcus Aureus. ACS Infect. Dis. 2020, 6 (10), 2771–2782. 10.1021/acsinfecdis.0c00503.

(11) Fellner, M.; Walsh, A.; Ahator, S. D.; Aftab, N.; Sutherland, B.; Tan, E. W.; Bakker, A. T.; Martin, N. I.; Stelt, M. van der; Lentz, C. S. Biochemical and Cellular Characterization of the Function of Fluorophosphonate-Binding Hydrolase H (FphH) in Staphylococcus Aureus Support a Role in Bacterial Stress Response. ACS Infect. Dis. 2023, 9 (11), 2119–2132. 10.1021/acsinfecdis.3c00246.

(12) Pouderoyen, G. van; Eggert, T.; Jaeger, K.-E.; Dijkstra, B. W. THE CRYSTAL STRUCTURE OF BACILLUS SUBTILIS LIPASE: A MINIMAL ALPHA/BETA HYDROLASE ENZYME. 2001. 10.2210/pdb1i6w/pdb.

(13) Kovacic, F.; Babic, N.; Krauss, U.; Jaeger, K.-E. Aerobic Utilization of Hydrocarbons, Oils, and Lipids; Rojo, F., Ed.; 2019; pp 255–289. 10.1007/978-3-319-50418-6_39.

(14) Khan, F. I.; Lan, D.; Durrani, R.; Huan, W.; Zhao, Z.; Wang, Y. The Lid Domain in Lipases: Structural and Functional Determinant of Enzymatic Properties. Front. Bioeng. Biotechnol. 2017, 5, 16. 10.3389/fbioe.2017.00016.

